## Supplement S3: Concentrations, uptake and radial patterns in optimized flow networks. for "On biological flow networks: Antagonism between hydrodynamic and metabolic stimuli as driver of topological transitions"

### S3 Appendix: Concentrations, uptake and radial patterns in optimized flow networks

In the following section we present additional material on the systematic parameter scan of linkwise demand-supply adaptation, as described in the results section 2. The diagrams depict the nodal concentrations, uptake effectiveness and radial distributions for the selected $\sigma_0$, $\beta^*$, $\alpha_0$, $\alpha_1$ variations depicted in Fig. 3 and 4. The $\chi$ notation depicts the relative distance of the respective vertex, or link center of the nearest source node, i.e. $\chi = 0$ corresponds to a source node while $\chi = 1$ describes sink-sided positions.

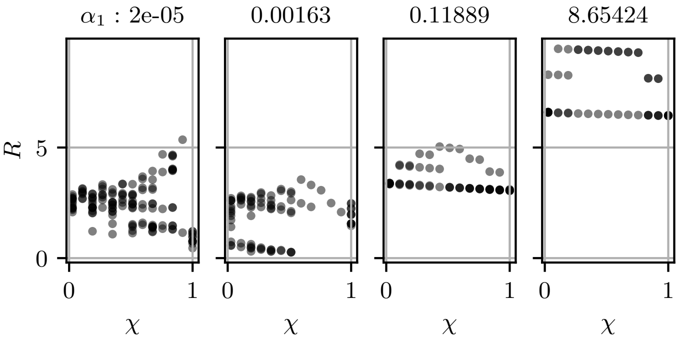


**(a)**

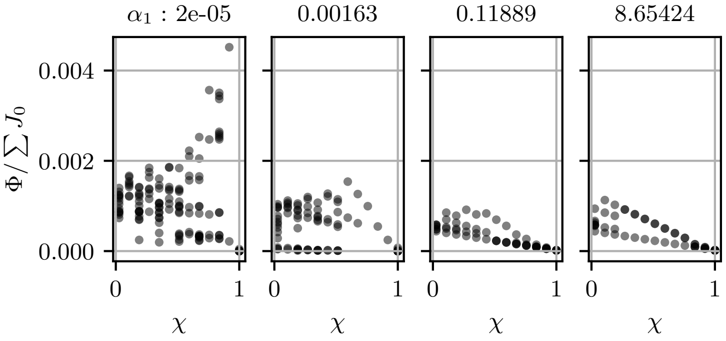


**(b)**

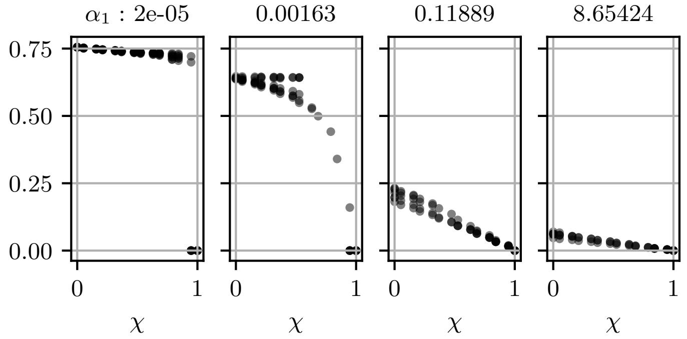


**(c)**

**Fig 1.** Radii, effective uptake and concentration profiles from source ($\chi = 0$) to sink ($\chi = 1$) for high demand and low absorption rates $\sigma_0 = 10^0$, $\beta^* = 10^{-3}$, $\alpha_0 = 4.5 \cdot 10^{-5}$, referring to the linkwise demand-supply scenario in hexagonal grids.

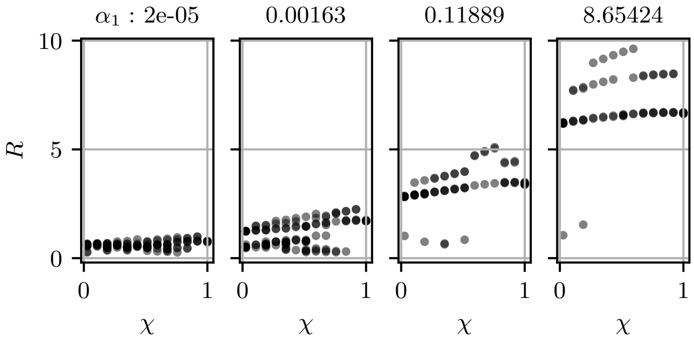


**(a)**

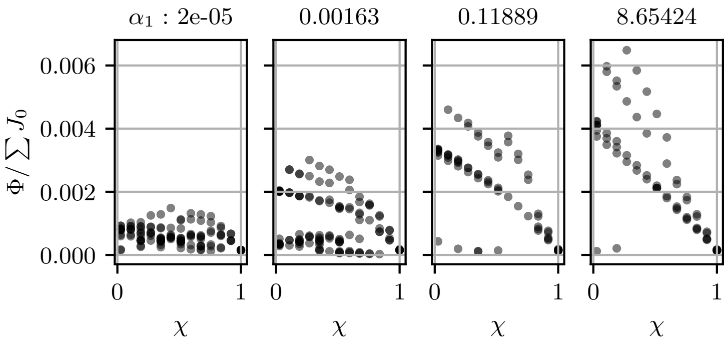


**(b)**

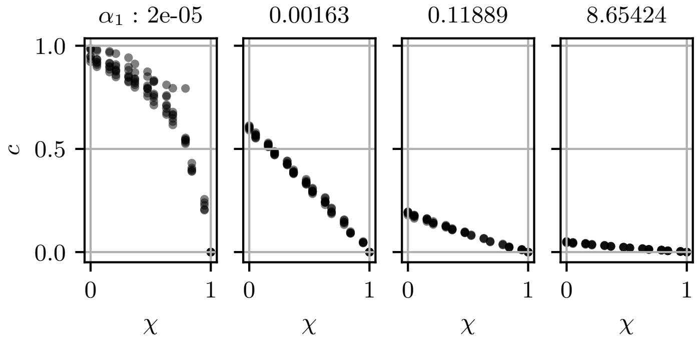


**(c)**

**Fig 2.** Radii, effective uptake and concentration profiles from source ($\chi = 0$) to sink ($\chi = 1$) for mediocre demand and mediocre absorption rates $\sigma_0 = 10^{-1}$, $\beta^* = 10^{-2}$, $\alpha_0 = 4.5 \cdot 10^{-5}$, referring to the linkwise demand-supply scenario in hexagonal grids.

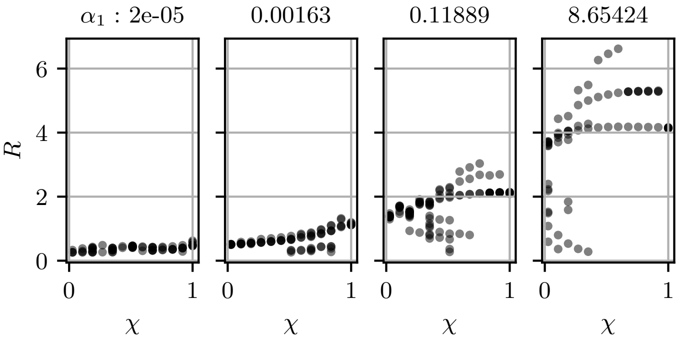


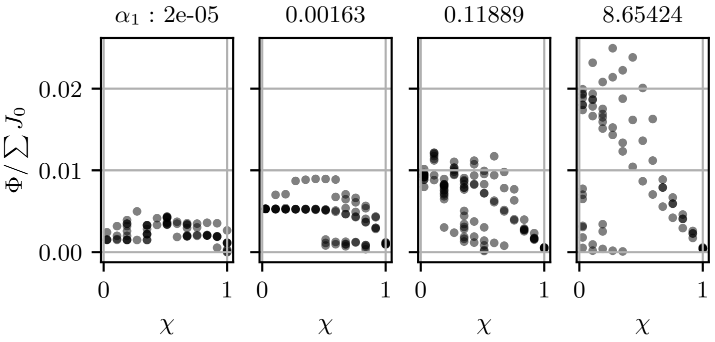


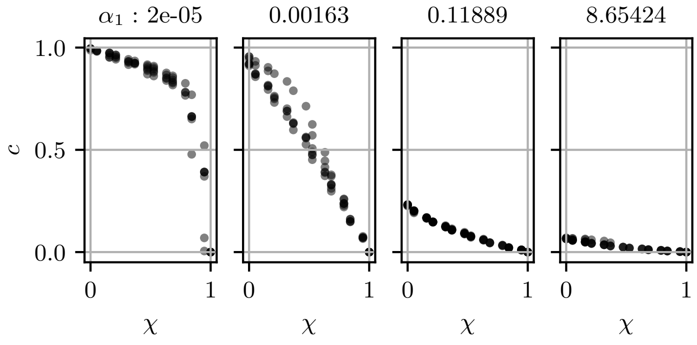


**(a)**

**(b)**

**(c)**

**Fig 3.** Radii, effective uptake and concentration profiles from source ($\chi = 0$) to sink ($\chi = 1$) for low demand and high absorption rates $\sigma_0 = 10^{-2}$, $\beta^* = 10^{-1}$, $\alpha_0 = 8 \cdot 10^{-8}$, referring to the linkwise demand-supply scenario in hexagonal grids.
