## Supplement S2: Demand-supply based adaptation algorithm. for "On biological flow networks: Antagonism between hydrodynamic and metabolic stimuli as driver of topological transitions"

### S2 Appendix: Demand-supply based adaptation algorithm

Now we formulate a dynamical system which shall describe the adaptation of a vessel network in order to minimize the metabolic cost function $\Gamma$:

$$\Gamma = S\left(\boldsymbol{\Phi}, \boldsymbol{\Phi}_0\right) + \sum_e \left(\alpha_1 \frac{f_e^2}{K_e} + \alpha_0 K_e^\gamma\right) \tag{1}$$

Following the gradient-descent approach in order reach Lyapunov stable states we formulate the equations of motions as,

$$\partial_t r_i = -\chi \partial_{r_i} \Gamma \tag{2}$$

$$= -\chi \sum_j \partial_{\Phi_j} S\left(\boldsymbol{\Phi}, \boldsymbol{\Phi}_0\right) \partial_{r_j} \Phi + \partial_{r_i} \sum_e \left(\alpha_1 \frac{f_e^2}{K_e} + \alpha_0 K_e^\gamma\right) \tag{3}$$

with an arbitrary $\chi > 0$ to ensure $\partial_t \Gamma \leq 0$. The last term in Eq. (3), concerning the derivatives of the dissipation-volume costs, has been discussed in previous studies , e.g. [1]. Subsequently we proceed with the metabolic uptake problem. First, we rewrite the uptake term $\Phi_k$ for an arbitrary link $k$ as

$$\Phi_k = \frac{q_k}{2}\left(\bar{c}_{\alpha(k)} G_k + \bar{c}_{\omega(k)} H_k\right) \tag{4}$$

with sub-functions defined as,

$$G_k = \left(x_k \coth\left(\frac{x_k}{2}\right) - \frac{x_k \mathrm{e}^{\frac{Pe_k}{2}}}{\sinh\left(\frac{x_k}{2}\right)} + Pe_k\right) \tag{5}$$

$$H_k = \left(x_k \coth\left(\frac{x_k}{2}\right) - \frac{x_k \mathrm{e}^{-\frac{Pe_k}{2}}}{\sinh\left(\frac{x_k}{2}\right)} - Pe_k\right) \tag{6}$$

and partial derivatives (using notation $\partial_{r_j}(\cdot) = \partial_j(\cdot)$) are calcuated as

$$\partial_j \Phi_k = \partial_j \frac{q_k}{2}\left(\bar{c}_{\alpha(k)} G_k + \bar{c}_{\omega(k)} H_k\right) + \frac{q_k}{2}\left(\partial_j \bar{c}_{\alpha(k)} G_k + \partial_j \bar{c}_{\omega(k)} H_k\right) + \frac{q_k}{2}\left(\bar{c}_{\alpha(k)} \partial_j G_k + \bar{c}_{\omega(k)} \partial_j H_k\right) \tag{7}$$

$$\partial_j q_k = 2\pi R_k \delta_{jk} \tag{8}$$

$$\partial_j c_i = \partial_j\left(\boldsymbol{e}_i^T \cdot \boldsymbol{c}\right) = \boldsymbol{e}_i^T \cdot \partial_j\left(\boldsymbol{M}^\dagger \boldsymbol{J}\right) = -\boldsymbol{e}_i^T \cdot \left(\boldsymbol{M}^\dagger \partial_j \boldsymbol{M} \boldsymbol{M}^\dagger \boldsymbol{J}\right) = -\boldsymbol{e}_i^T \cdot \left(\boldsymbol{M}^\dagger \partial_j \boldsymbol{M} \boldsymbol{c}\right) \tag{9}$$

Subsequently we can calulate the derivates of $\boldsymbol{M}$ and the sub-components as

$$\begin{aligned}
\partial_j M_{ij} = &\sum_k \partial_j \frac{q_k}{2}\left[B_{ik} Pe_k + |B_{ik}| x_k \coth\left(\frac{x_k}{2}\right)\right]\delta_{ij} \\
&+ \sum_k \frac{q_k}{2}\left(B_{ik} + |B_{ik}|\left\{\frac{\coth\left(\frac{x_k}{2}\right)}{x_k} - \left[\frac{\coth\left(\frac{x_k}{2}\right)}{\cosh\left(\frac{x_k}{2}\right)}\right]^2\right\}\right)\partial_j Pe_k \delta_{ij} \\
&- \sum_{k \in out(i)}\left\{\partial_j \frac{q_k}{2}\frac{x_k \mathrm{e}^{-\frac{Pe_k}{2}}}{\sinh\left(\frac{x_k}{2}\right)} + \frac{q_k}{2}\frac{\mathrm{e}^{-\frac{Pe_k}{2}}}{\sinh\left(\frac{x_k}{2}\right)}\left[\frac{Pe_k}{x_k} - \frac{x_k}{2} - \frac{Pe_k}{2}\coth\left(\frac{x_k}{2}\right)\right]\partial_j Pe_k\right\}\delta_{\omega(k),j} \\
&- \sum_{k \in in(i)}\left\{\partial_j \frac{q_k}{2}\frac{x_k \mathrm{e}^{-\frac{Pe_k}{2}}}{\sinh\left(\frac{x_k}{2}\right)} + \frac{q_k}{2}\frac{\mathrm{e}^{-\frac{Pe_k}{2}}}{\sinh\left(\frac{x_k}{2}\right)}\left[\frac{Pe_k}{x_k} + \frac{x_k}{2} - \frac{Pe_k}{2}\coth\left(\frac{x_k}{2}\right)\right]\partial_j Pe_k\right\}\delta_{\alpha(k),j}
\end{aligned} \tag{10}$$

$$\partial_j G_k = \frac{\partial_j Pe_k}{\sinh\left(\frac{x_k}{2}\right)}\left\{\frac{Pe_k}{x_k}\left[\cosh\left(\frac{x_k}{2}\right) - \mathrm{e}^{\frac{Pe_k}{2}}\right] + \sinh\left(\frac{x_k}{2}\right) - \frac{Pe_k}{2\sinh\left(\frac{x_k}{2}\right)} + \mathrm{e}^{\frac{Pe_k}{2}}\left[\frac{Pe_k}{2}\coth\left(\frac{x_k}{2}\right) - \frac{x_k}{2}\right]\right\} \tag{11}$$

$$\partial_j H_k = \frac{\partial_j Pe_k}{\sinh\left(\frac{x_k}{2}\right)}\left\{\frac{Pe_k}{x_k}\left[\cosh\left(\frac{x_k}{2}\right) - \mathrm{e}^{-\frac{Pe_k}{2}}\right] - \sinh\left(\frac{x_k}{2}\right) - \frac{Pe_k}{2\sinh\left(\frac{x_k}{2}\right)} + \mathrm{e}^{-\frac{Pe_k}{2}}\left[\frac{Pe_k}{2}\coth\left(\frac{x_k}{2}\right) + \frac{x_k}{2}\right]\right\} \tag{12}$$

Finally we calculate the derivates of the local Peclet numbers as

$$\partial_j Pe_k = \frac{L_k}{D}\partial_j \bar{u}_k \text{ with } \partial_j \bar{u}_k = \partial_j\left(\frac{R_k^2 \Delta p_k}{8\eta L_k}\right) = \frac{1}{8\eta L_k}\left(2\delta_{jk} R_k \Delta p_k + R_k^2 \partial_j \Delta p_k\right) \tag{13}$$

We compute the pressure derivatives from the Kirchhoff solutions as

$$\partial_j \Delta p_k = \partial_j \left[ \boldsymbol{e}_k^T \boldsymbol{B}^T \left( \boldsymbol{B}\boldsymbol{C}\boldsymbol{B}^T \right)^{\dagger} \boldsymbol{s} \right] \tag{14}$$

$$= -\boldsymbol{e}_k^T \boldsymbol{B}^T \left[ \boldsymbol{B}\boldsymbol{C}\boldsymbol{B}^T \right]^{\dagger} \boldsymbol{B} \partial_j \boldsymbol{C} \Delta \boldsymbol{p} \tag{15}$$

$$= -\boldsymbol{e}_k^T \boldsymbol{B}^T \left[ \boldsymbol{B}\boldsymbol{C}\boldsymbol{B}^T \right]^{\dagger} \boldsymbol{B} \left[ \frac{4C_j}{R_j} \left( \boldsymbol{e}_j \otimes \boldsymbol{e}_j \right) \right] \Delta \boldsymbol{p} \tag{16}$$

$$= -\frac{4C_j \Delta p_j}{R_j} \left( \boldsymbol{B}^T \left[ \boldsymbol{B}\boldsymbol{C}\boldsymbol{B}^T \right]^{\dagger} \boldsymbol{B} \right)_{jk} \tag{17}$$

which closes the system derivatives for $\partial_t r_i$ and allows us to evaluate the flow landscape as well as the occuring gradients accordingly. For numerical purposes, one is remined to carefully evaluate Peclet number dependend exponential and hyperbolical functions as overflow errors are likely to occur.
