## Supplement S1: Metabolite transport in arbitrary Kirchhoff networks. for "On biological flow networks: Antagonism between hydrodynamic and metabolic stimuli as driver of topological transitions"

### S1 Appendix: Metabolite transport in arbitrary Kirchhoff networks

In this section we shall give a detailed run-down of the derivation of the equations systems and solutions in section 1. The continuity equation was reading in the non-dimensional form as

$$\bar{c}\left(z^*\right) = X_0 \mathrm{e}^{\lambda_+ z^*} + X_1 \mathrm{e}^{\lambda_- z^*} \tag{1}$$

$$\text{with } \lambda_\pm = \frac{1}{2}\left(Pe \pm \sqrt{Pe^2 + \beta^*}\right) \tag{2}$$

with channels boundaries $\bar{c}\left(z=0\right) = \bar{c}_0$, $\bar{c}\left(z=L\right) = \bar{c}_L$ and $X_0 = \frac{\bar{c}_L - \bar{c}_0 \mathrm{e}^{a_1}}{\mathrm{e}^{a_0} - \mathrm{e}^{a_1}}$, $X_1 = \frac{\bar{c}_0 \mathrm{e}^{a_0} - \bar{c}_L}{\mathrm{e}^{a_0} - \mathrm{e}^{a_1}}$. In order to calculate the solute flux and the corresponding equations system for a complex network one writes (using $x_e = \sqrt{Pe_e^2 + \beta_e^*}$)

$$I_k\left(z\right) = A_k\left(\bar{u}_k \bar{c}_k\left(z\right) - D\partial_z \bar{c}_k\left(z\right)\right) = \frac{A_k D}{L}\left(Pe_k \bar{c}_k\left(z\right) - \partial_{z^*}\bar{c}_k\left(z\right)\right) \tag{3}$$

$$\Rightarrow I_k\left(0\right) = I_{\alpha(k)} = \frac{A_k D}{2L}\left\{\bar{c}_{\alpha(k)}\left[Pe_k + x_k \coth\left(\frac{x_k}{2}\right)\right] - \bar{c}_{\omega(k)} \frac{x_k \mathrm{e}^{-\frac{Pe_k}{2}}}{\sinh\left(\frac{x_k}{2}\right)}\right\} \tag{4}$$

$$\Rightarrow I_k\left(L\right) = I_{\omega(k)} = \frac{A_k D}{2L}\left\{\bar{c}_{\omega(k)}\left[Pe_k - x_k \coth\left(\frac{x_k}{2}\right)\right] + \bar{c}_{\alpha(k)} \frac{x_k \mathrm{e}^{\frac{Pe_k}{2}}}{\sinh\left(\frac{x_k}{2}\right)}\right\} \tag{5}$$

Now we utilize the the boundary conditions for solute flux on every node as, using $\frac{A_k D}{L} = q_k$ as an abbreviation, so that we write for in- and outflux of solute of a vertex:

$$J_i = \sum_k B_{ik} I_k = \sum_{k \in out(i)} I_{\alpha(k)} - \sum_{k \in in(i)} I_{\omega(k)} \tag{6}$$

$$= \sum_k q_k \left(B_{ik} Pe_k + |B_{ik}| x_k \coth\left(\frac{x_k}{2}\right)\right)\bar{c}_i$$

$$- \sum_{k \in out(i)} q_k \frac{x_k \mathrm{e}^{-\frac{Pe_k}{2}}}{2\sinh\left(\frac{x_k}{2}\right)} \bar{c}_{\omega(k)} - \sum_{k \in in(i)} q_k \frac{x_k \mathrm{e}^{\frac{Pe_k}{2}}}{2\sinh\left(\frac{x_k}{2}\right)} \bar{c}_{\alpha(k)} \tag{7}$$

We may rewrite this equation system according to [1] in vectorial form,

$$\boldsymbol{M} \cdot \boldsymbol{c} = \boldsymbol{J} \tag{8}$$

which also allows us to efficiently solve for $\boldsymbol{c}$ for any given $\boldsymbol{J}$ with

$$M_{ij} = \sum_k \frac{q_k}{2}\left[B_{ik}Pe_k + |B_{ik}|x_k \coth\left(\frac{x_k}{2}\right)\right]\delta_{ij} - \sum_{k\in out(i)} q_k \frac{x_k \mathrm{e}^{-\frac{Pe_k}{2}}}{2\sinh\left(\frac{x_k}{2}\right)}\delta_{\omega(k),j} - \sum_{k\in in(i)} q_k \frac{x_k \mathrm{e}^{\frac{Pe_k}{2}}}{2\sinh\left(\frac{x_k}{2}\right)}\delta_{\alpha(k),j} \tag{9}$$

As we generally utilize mixed boundary conditions with $c_n = 0$ on the outflux periphery and $J_n > 0$ on the influx periphery of the network we can calculate the complementary missing values and subsequently calculate the linkwise metabolite absorption as,

$$\Phi = \sum_i J_i \tag{10}$$

$$= \sum_k \frac{q_k}{2}\left\{\bar{c}_{\alpha(k)}\left[x_k \coth\left(\frac{x_k}{2}\right) - \frac{x_k \mathrm{e}^{\frac{Pe_k}{2}}}{\sinh\left(\frac{x_k}{2}\right)} + Pe_k\right] + \bar{c}_{\omega(k)}\left[x_k \coth\left(\frac{x_k}{2}\right) - \frac{x_k \mathrm{e}^{-\frac{Pe_k}{2}}}{\sinh\left(\frac{x_k}{2}\right)} - Pe_k\right]\right\} \tag{11}$$

$$\Rightarrow \Phi_k = \frac{q_k}{2}\left\{\bar{c}_{\alpha(k)}\left[x_k \coth\left(\frac{x_k}{2}\right) - \frac{x_k \mathrm{e}^{\frac{Pe_k}{2}}}{\sinh\left(\frac{x_k}{2}\right)} + Pe_k\right] + \bar{c}_{\omega(k)}\left[x_k \coth\left(\frac{x_k}{2}\right) - \frac{x_k \mathrm{e}^{-\frac{Pe_k}{2}}}{\sinh\left(\frac{x_k}{2}\right)} - Pe_k\right]\right\} \tag{12}$$

In Fig. 1 we display an examplary solution for a single channel with absorbing boundary. One finds the concentrations profile in general to become linear in the case of small Peclet numbers, corresponding to the maximal possible decline in solute flux, see 1a. Further to get a quantitative and qualitative perspective on the metabolite uptake in this system one may consider the effective uptake, see 1b. As depicted one finds the uptake to vary drastically with the Peclet numbers as well as the effective absorption rate. Note that increased Peclet numbers generally correspond to a decreased uptake while entering the diffusive regime for $Pe \to 0$ displays clear saturation behavior in dependence of $\beta^*$ .

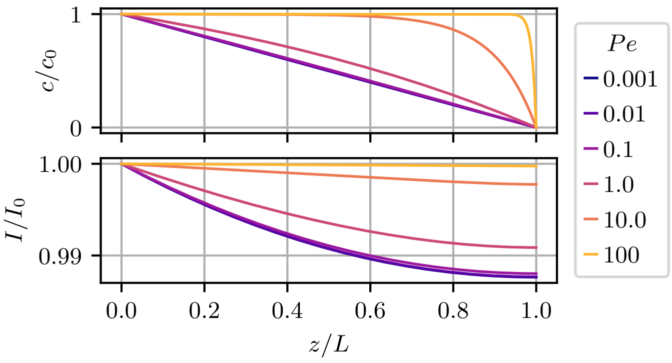


**(a)**

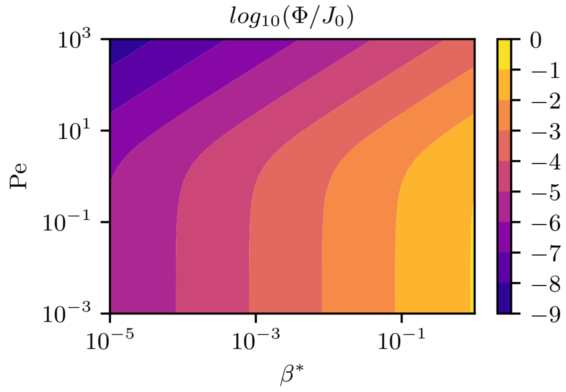


**(b)**

**Fig 1.** Single channel solutions with absorbing boundary: (a) Concentration and solute flux profiles for $\beta^* = 10^{-2}$ (b) Effective metabolite uptake
